# Combined effects of 12-week yoga warm-up on athletic performance in male high school track and field athletes

**DOI:** 10.1101/2022.04.14.488191

**Authors:** Danyang Wei

**Affiliations:** Department of Sports Science, Xi’an Physical Education University, Xian, China

**Keywords:** performance, sEMG, yoga, stretching

## Abstract

**Background:** Practicing yoga could improve balance and flexibility, but its positive significance as a long-term warm-up for formal training was uncertain. We hypothesized that practicing yoga during warm-up might positively affect balance, flexibility, and speed performance in male high school track and field athletes.

**Methods:** Over a 12-week period, athletes in a yoga group (YG) (n=10) practiced yoga for 15 min 4 times a week during warm-up, while athletes in a dynamic stretching group (DSG) (n=10) practiced 15 min of dynamic stretching. Except for the warm-up activities, the training content of the two groups of students was the same. we tested performance indicators immediately before and after the intervention, including lower extremity flexibility test (right hip active flexion range), lower extremity balance test [using surface electromyography (sEMG) to measure right leg tibialis anterior (TA) activation during one-leg stance (OLS) with eyes closed], and speed performance test (100-meter and 800-meter tests).

**Results:** We performed between-group and within-group comparisons for indicators of two groups by using SPSS (version 26.0). Within-group comparisons showed a significant improvement in flexibility (P=0.002) and balance (P=0.003) in YG, but no significant change in DSG, after the 12-week intervention. In addition, speed performance of both YG (100m, P=0.026; 800m, P=0.045) and DSG (100m, P=0.029; 800m, P=0.006) was significantly improved. Between-group comparison showed that YG had a significant advantage in 800m (P=0.045) and flexibility (P=0.031).

**Conclusions:** These data suggested that practicing yoga as a long-term warm-up could help male high school track and field athletes improve lower body flexibility and 800m speed. In addition, yoga had a certain positive effect on balance, but it was not significant overall.

## 1 Introduction

How to design a warm-up has been controversial. The prevailing view was that dynamic stretching as well as static stretching should be used as a pre-training warm-up [1, 2], and proponents of this view believed that this type of training could increase range of motion, reduce muscle resistance, and improve athletic performance [3, 4]. But now this mainstream view is being challenged.

Static stretching was used in warm-up to increase the range of motion [5]. However, now studies have shown that repetitive and sustained static stretching may impair muscle strength and speed performance, also increase the risk of sports injury during subsequent exercise due to joint instability [6, 7, 8, 9]. Due to the negative effects of static stretch, dynamic stretching was used more in warm-up activities, which showed positive effects on vertical jump height, maximal muscle strength, and sprint speed [10, 11, 12]. However, there were also some studies showing that dynamic stretching had no positive effect on these performances [13, 14].

We found some studies showing that stretching had a positive effect on athletic performance, while others found the opposite. We speculated the reason was that there were differences in the design of warm-up activities by different researchers, including different forms of stretching combinations, movement design, intensity and duration [15, 16, 17], which would affect the effectiveness of warm-up activities. In order to design more effective warm-up activities, researchers have conducted various explorations. Lykesas et al. [18] designed a warm-up routine using Greek traditional dance, and found no difference in flexibility performance and better performance in agility tests compared to dynamic stretching in immediate response. Chen et al. [19] added a vibrating foam roller to a traditional warm-up routine and found that this combination significantly improved performance on the hexagon test and the frequency speed of kick test (FSKT). Pojskić et al. [20] used prolonged intermittent low-intensity isometric exercise as a warm-up program and found that when 30 % body weight resistance was applied, the improvements of speed and agility were similar to dynamic stretching.

In the exploration of warm-up activity design, yoga was introduced [21, 22], as practicing yoga could increase heart rate and body temperature [23] and stretch 97% of the muscles in the body [24]. Yoga practice could smooth the contraction of skeletal muscles [25], which allowed joints to be stretched to a greater extent [26]. In addition, yoga always choreographed asanas in sequence, and this dynamic pattern helped improve movement coordination and muscle recruitment efficiency [27]. By practicing yoga, trainers could be more effective at developing flexibility [28, 29] and balance [30]. However, we found that these studies did not comprehensively analyze flexibility, balance, and athletic performance.

The purpose of this paper was to address these limitations and to analyze the combined effects of practicing yoga as a long-term warm-up, including: flexibility, balance, and speed performance on high school male track and field athletes,

## 2 Method

### 2.1 Participant

The participants in this experiment were high school track and field athletes from Xi’an Soaring Sports Club, a total of 20 young males. The average age of these participants was 17 years old, and their training experience did not exceed 6 months. All players were free from any current or previous injuries and were not taking drugs that affected balance. In addition they had no prior experience with yoga and were able to make sure to train on time during the experiment. We randomly assigned those athletes to a yoga group (YG, N=10) and a dynamic stretching group (DSG, N=10), which was shown in Table 1.

**Table 1:** Participant Demographics by Group.

|  | DSG | YG | P |
| --- | --- | --- | --- |
| Height(cm) | 175.5 | 176.9±4.2 | 0.439 |
| Weight (kg) | 65.0±2.7 | 63.9±5.0 | 0.547 |
| Age (years) | 16.8±0.6 | 17.0±0.5 | 0.433 |
| Training experience(months) | 3.7±0.8 | 4.2±1.1 | 0.274 |
| BMI | 21.1±0.6 | 20.4±1.2 | 0.124 |
| Attendance over 12-week intervention | 45.8±2.0 | 46.6±2.0 | 0.376 |

### 2.2 Experimental Procedures

Over a 12-week period, athletes of both groups completed exercise training sessions 4 times a week for a total of 48 sessions. YG athletes practiced yoga for 15 min after 10 min of jogging as a warm-up training, while DSG athletes practiced dynamic stretching for 15 min. The training content of the two groups was the same except for the warm-up training, and these athletes were asked not to participate in additional physical activity. Flexibility, balance, and speed performance were measured before and after the experiment.

Yoga Intervention is designed by certified yoga instructors, which was a simple but progressive 12-week program. Based on Suryanamaskar, this yoga program added several asanas focused on lower limb flexibility and balance, and all these asanas were performed in series. YG athletes needed to complete 5 times yoga procedure within 15 minutes, while stretching movements in DSG were mainly in the marching style, followed by some standing and sitting stretches.

### 2.3 Measurements

Flexibility: We measured the athlete’s right hip active flexion range [31] by SYJDC200 angle ruler and averaged three measurements. During the measurement, the athlete lied flat on the mat, one tester immobilizing the pelvis, and then the athlete straightened the right leg and raised it up to the maximum angle.

Balance: We chose the one-leg standing test (OLS) [32] with subjects’ eyes closed, and then recorded the surface electromyography (sEMG) signal of the right leg tibialis anterior (TA) during this period. Before the test, we used a razor to shave off the body hair of the test site and wiped it with alcohol. According to the suggestion of SENIAM [33], the electrode was pasted on the corresponding part from TA. We used a 6-channel sEMG sensor, setting sampling frequency to 1000Hz, and the bandpass filter frequency to 20-500Hz. Before the start of the experiment, we measured the root mean square (RMS) of the subjects’ TA during the maximum voluntary isometric contraction (MVIC) in dorsiflexion for 5s. This test was repeated twice with 5 min intervals, and the maximum value was taken as the record. During the OLS test, subjects were briefly destabilized for the first few seconds after closing their eyes. To alleviate this interference, we recorded sEMG 5 s after the start of the test for a total of 10s, repeated twice, with 5 min intervals between each measurement. We calculated the RMS average of the two measurements from the TA, and divided it by the RMS during maximal voluntary isometric contractions during dorsiflexion, using %RMS for further discussion.

Speed performance: We measured the time taken by the athletes for the 100m and 800m tests twice and recorded the best time.

### 2.4 Data Analysis

Before all hypothesis testing, independent samples t-tests were performed using SPSS 26 for Windows (SPSS, Inc., Chicago, IL, USA) to examine whether YG and DSG were significantly different in demographics, flexibility, balance, and speed performance. We could not use ANCOVA as the test showed that the regression lines for the 100m and 800m speed data for the two groups were not parallel. The T test showed that the data of the two groups were basically the same at baseline, so we chose independent samples T test and paired samples T test for intra- and between-group comparisons to analyze changes in two groups. Significance (*) was set at the alpha level of P = 0.05, and data given represents the mean ± SD unless stated otherwise.

## 3 Results

Table2 and Table3 were shown the information on the between-group and within-group comparison respectively. We could see that the indicators of two groups were basically the same at the baseline, as the T test results showed that there was no significant difference. Since these athletes were novices, their speed performance improved significantly through 12 weeks of training. The intra-group comparison showed that the time of DSG at 100m and 800m tests was shortened by 0.43s (P<0.05) and 5.4s (P<0.01) respectively, while YG was shortened by 0.37s (P<0.05) and 7.0s (P<0.05) in these two items. The between-group comparison showed that the speed performance of YG at 800m was better than that of DSG after intervention (P<0.05).

**Table 2:**
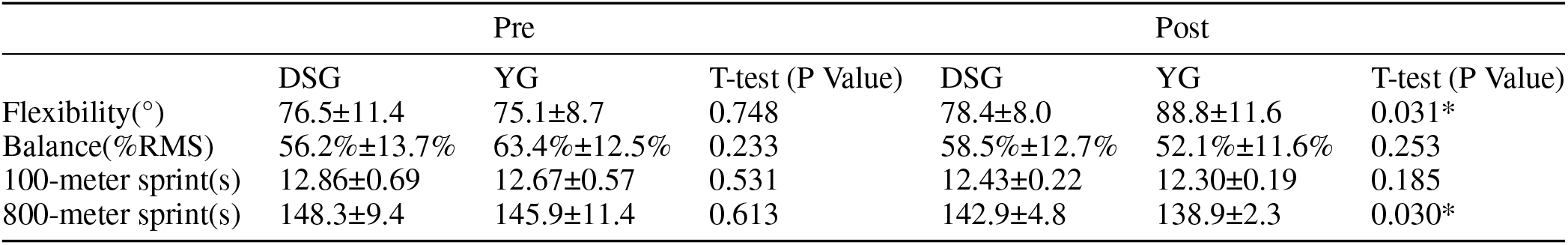
Comparison of flexibility, balance and speed performance between YG and DSG.

**Table 3:** Flexibility, balance, and speed performance changes in YG and DSG.

|  | DSG |  | YG |  |
| --- | --- | --- | --- | --- |
|  | Post-pre change | T-test(P Value) | Post-pre change | T-test(P Value) |
| Flexibility($^{\circ}$) | 1.9±4.6 | 0.235 | 13.7±10.4 | 0.002* |
| Balance(%RMS) | 2.3%±5.5% | 0.221 | -11.3%±8.7% | 0.003* |
| 100-meter sprint(s) | -0.43±0.52 | 0.029* | -0.37±0.45 | 0.027* |
| 800-meter sprint(s) | -5.4±4.8 | 0.008* | -7.0±9.4 | 0.043* |

The flexibility and balance indicators in two groups improved significantly after the intervention. The active flexion angle of the right hip joint increased by 13.7° (P < 0.01), and the sEMG signals of the right TA during balance maintenance decreased by 11.3%RMS (P < 0.01), while DSG hardly changed on these two indicators. When comparing the data between groups after the intervention, YG showed a significant advantage in flexibility (P<0.05), but no such advantage was observed in balance. The reason might be that there was a certain degree of individual differences in the balance indicators of two groups at baseline.

## 4 Discussion

In this study, both YG and DSG participated in a complete training program, including warm-up exercise, formal training, and finishing exercise. Athletes in both groups were novices and their training content was the same except for warm-up activities. Therefore, we expected an improvement in flexibility, balance, and speed performance in both groups, while the experimental results were in line with expectations to a certain extent.

Our findings showed that practicing yoga as a long-term warm-up improved flexibility in these male athletes, whereas dynamic stretching did not. Furthermore, although not significant overall, yoga practice improved the athlete’s balance. In terms of speed performance, both YG and DSG have improved, but compared to DSG (−3.6%), YG (−4.8%) has a greater speed improvement at 800m, and the difference is significant.

### 4.1 Effects of practicing yoga as a long-term warm-up on flexibility

Previous studies found that yoga appeared to be more effective than dynamic stretching for improving flexibility in both immediate [34] and long-term [35] effects, but these studies did not practice yoga during warm-up, whereas this experiment demonstrated that yoga as a warm-up still valid. There was no change in DSG flexibility, which we speculated was due to the lack of flexibility exercises in the training programs of these athletes. A previous study showed that athletes improved strength but decreased shoulder flexibility after weightlifting [36].

We analyzed that yoga could effectively improve flexibility from two aspects. First, the practice of yoga was not likely to induce stretch reflexes [37]. The practice of yoga involved short active contractions of the stretched muscle before stretching, which may reduce the sensitivity of muscle spindles and allow for greater stretches. Second, the training content of yoga was highly structured, which seemed to be more efficient. The asanas were carried out in series during the practice, and it only took a short time to change from one movement to the next, allowing more content to be arranged in a limited time.

### 4.2 Effects of practicing yoga as a long-term warm-up on balance

This study found that practicing yoga during warm-up appeared to be more effective than dynamic stretching for improving balance performance. The effect of dynamic stretching on balance has been controversial [38, 39, 40, 41], while yoga has been shown to be positive for balance in different populations [42, 43, 44, 45]. The training of ankle flexibility and stability in yoga asana practice accounted for a large proportion [46, 47], which made it important for running, because ankle injuries accounted for 1/3 of running injuries [48] and ankle stability also affected the efficient in running [49].

We used sEMG to demonstrate the positive effects of yoga on balance, which was rare in previous studies. Previous studies showed that sEMG was able to accurately measure muscle activity through bioelectrical changes [50, 51, 52]. TA played an important role in the stabilization of OLS [53], and the stability of the sEMG signal amplitude during the process of maintaining balance meant the improvement of balance efficiency. So, the mechanism was using less muscle activation to accomplish the same task [54]. We noticed that YG significantly decreased the percentage of tibialis anterior muscle EMG when maintaining balance, which proved the positive effect of yoga on balance on a physiological level.

### 4.3 Effects of practicing yoga as a long-term warm-up on speed performance

A previous study showed that after 10 weeks of yoga intervention, the yoga group presented an advantage in flexibility compared to the control group, but there was no significant difference in 30m sprint performance [55]. Our conclusion was similar to this study, as YG also did not show an advantage in the speed performance of 100m. But unlike previous studies, YG achieved a bigger improvement at 800m. We speculated that the reason for this result was that as flexibility and balance improve, the economy of running improved.

Practicing yoga improved flexibility, which reduced active stiffness and improved muscle performance in stretch shortening cycle (SSC) [56]. By increasing the elastic potential energy stored in the muscles during the SSC, athletes could generate greater force [57, 58]. In addition, the viscosity of the muscle decreased, reducing the energy loss during SSC [59]. Improvements in balance could reduce the proportion of stabilizing muscles involved in balance tasks [60], which helped reduce erratic movements [61] and enhanced the synergistic contraction of agonist and antagonist muscles [62]. Ultimately these factors all increased the efficiency of the exercise.

Practicing yoga as a long-term warm-up improved the efficiency of exercise. Compared to DSG, YG showed a significant advantage in speed performance at 800m, but no difference at 100m. We speculated that the effect of sport economy on speed performance needed more time to manifest, because it took a lot longer to complete 800m than 100m, which allowed the movement economy to accumulate.

The focus of this study was to discuss the combined effects of yoga as a warm-up activity on high school track and field athletes, so we did not design additional groups to differentiate between the effects of flexibility and balance on exercise economy, which prevented us from discussing the impact of these two qualities on speed performance separately.

## 5 Conclusion

After 12 weeks of training, male track and field athletes who practiced yoga as a warm-up has significantly improved flexibility compared to dynamic stretching. Additionally, although not significantly overall, yoga practice improved the athlete’s balance. Finally, yoga practice boosted athlete’s 800m speed, which we analyzed was because yoga improved running economy. Therefore, we recommended that novice male track and field athletes practiced yoga sequences in general warm-up activities, which could help athletes improve balance, flexibility, and middle-distance running performance.

## 6 Financial support and sponsorship

Nil

## 7 Conflicts of interest

There are no conflicts of interest.

## 8 Acknowledgment

We would like to thank the coaches and participants of the study for their time.

## Notes

### Competing Interest Statement

The authors have declared no competing interest.

